# Unicellular cyanobacteria rely on sodium energetics to fix N_2_

**DOI:** 10.1101/2024.03.08.584021

**Authors:** Si Tang, Xueyu Cheng, Yaqing Liu, Lu Liu, Dai Liu, Qi Yan, Jianming Zhu, Jin Zhou, Katrin Hammerschmidt, Zhonghua Cai

## Abstract

Diazotrophic cyanobacteria can thrive in combined nitrogen (N)-limited environments due to their ability to fix nitrogen gas (N_2_) from the atmosphere. Despite this, they occur in low abundance in N-limited coastal waters, which represents an ecological paradox^1–3^. One hypothesis is that this is partly due to elevated salinity (> 10 g/L NaCl), which inhibits cyanobacterial N_2_ fixation^2,3^. Here we show that N_2_ fixation in a unicellular coastal cyanobacterium is not inhibited but rather exclusively dependent on sodium (Na^+^) ions. In N-deficient environments, both N_2_ fixation and population growth were significantly inhibited at low NaCl concentrations (< 4 g/L). Additional experiments indicated that sodium energetics, rather than proton energetics, is necessary for N_2_ fixation, as Na^+^ deficiency resulted in insufficient ATP supply for N_2_ fixation. We show that this is due to the non-functioning Na^+^-coupled ATP synthase, which we found to be likely coupled to anaerobic rather than aerobic respiration. Sequence alignment analysis of the ion-coupling site of the ATP synthase revealed a high prevalence of Na^+^ energetics in cyanobacteria, with all unicellular N_2_ fixers capable of Na^+^ energetics. This suggests a critical role for sodium energetics in cyanobacteria. It also raises the possibility that sodium energetics is not as rare as thought, but that we may have underestimated the prevalence and importance of sodium energetics in other organisms. Finally, the low abundance of diazotrophic unicellular cyanobacteria in coastal waters may be due to insufficient NaCl levels to support N_2_-fixation during periods of growth-supporting high temperatures. This provides another perspective on the regulation of the oceanic N cycle that needs to be considered in times of global climate change. Changes in current patterns could lead to an overlap of periods optimal for N_2_ fixation and population growth, likely resulting in dense cyanobacterial blooms.

## Main

Organisms require combined nitrogen (N) for essential processes such as nucleic acid and protein synthesis. Despite the abundance of atmospheric nitrogen gas (N_2_), bioavailable N is often scarce in nature^4^. In aquatic ecosystems, cyanobacteria can partially meet the need for N by fixing N_2_, which impacts food chains, greenhouse gas sequestration, global climate change, and biogeochemical cycles^4,5^. This ability provides diazotrophic cyanobacteria with a competitive advantage over non-diazotrophic cyanobacteria and eukaryotic phytoplankton when environmental N is low. It enables them to exploit and dominate N-scarce environments worldwide, and can even lead to the development of dense cyanobacterial blooms when other nutrients are abundant^6^. Interestingly, unlike the frequently reported blooms of cyanobacterial N_2_-fixers in freshwater ecosystems, there are only a few reported blooming events in N-limited saline coastal waters, even though they have a clear ecological advantage^2,3^. The scarcity and low abundance of coastal N_2_-fixing cyanobacteria in an N-deficient environment is striking and presents an ecological paradox.

One commonly proposed explanation is that high salinity levels (> 10 g/L NaCl), a fundamental difference between coastal water and freshwater ecosystems, severely inhibit N_2_ fixation and population development of resident N_2_-fixers^2,3,7^. However, most of these studies refer to multicellular N_2_-fixing cyanobacteria and therefore much remains unknown about the NaCl tolerance of coastal unicellular N_2_-fixers, which have only recently been recognised for their significant contribution to the oceanic N cycle^4,8^.

Here we investigate the effect of NaCl on N_2_ fixation using *Cyanothece* sp. ATCC 51142 (hereafter *Cyanothece* sp.), a coastal N_2_-fixing unicellular cyanobacterium^9^. First, we investigate whether high salinities suppress unicellular cyanobacterial N_2_ fixation, as proposed for multicellular cyanobacteria. We then integrate physiological, transcriptomic, and enzymatic analyses to investigate the underlying mechanisms. Finally, through a protein sequence alignment study, we expand on our findings in the cyanobacteria phyla, and also discuss their role in the global N cycle.

Contrary to the accepted view that high salinity negatively affects coastal N_2_ fixation^2,3^, here we demonstrate not only that N_2_ fixation of *Cyanothece* sp. is NaCl-dependent, but rather that Na^+^ energetics is essential to provide ATP for N_2_ fixation. Sequence alignment analysis of the ion-coupling site of the ATP synthase revealed a surprisingly high prevalence of Na^+^ energetics capability – particularly in marine cyanobacteria. Thus, in N-deficient coastal areas, freshwater input from rivers and precipitation leads to low salinities during times with growth-favouring high temperatures and vice versa. This may explain the low rates of N_2_ fixation and the low abundance of cyanobacterial N_2_-fixers, resulting in N limitation in coastal seas. Overall, we provide an alternative perspective on the paradox of the scarcity of N_2_-fixing cyanobacteria in N-limited coastal waters. Moreover, we provide insights into the role of Na^+^ energetics in cyanobacterial metabolism, specifically in N_2_ fixation.

## Results

### N_2_ fixation depends on NaCl presence

We first carried out growth experiments of *Cyanothece* sp. cultured in artificial seawater medium (ASP2, 18 g/L NaCl) and freshwater medium (BG11) in the presence or absence of N. Under N-rich conditions, cells from ASP2 and BG11 showed similar population growth, as expected for a coastal isolate. Under N deficiency, population growth was only observed in ASP2 without N (hereafter ASP2-N), whereas cells barely grew in BG11 without N (hereafter BG11_0_) (Fig. 1a). Since the most prominent discrepancy between ASP2-N and BG11_0_ is the presence of NaCl, we recultured these non-growing cells in fresh BG11_0_ amended with 18 g/L NaCl (hereafter BG11_0_ (NaCl)), and NaCl reactivated population growth (Fig. 1b). Here N_2_ fixation is the prerequisite for cell growth and division, it is therefore confirmed that N_2_ fixation depends on NaCl. The effect of NaCl gradients on N_2_ fixation and growth was investigated in more detail. When N was replete, except for the three highest NaCl concentrations (32, 34, 36 g/L) (ANOVA, F_3,8_ = 22.04, *p* < 0.001), cells in the other treatments did not differ in growth from cells grown at 18 g/L, which is equivalent to the salinity of the marine medium ASP2 (Fig. 1c). In contrast, when N was depleted, NaCl was required for N_2_ fixation and population growth. Cell proliferation was almost completely inhibited when NaCl was less than 4 g/L and maximised at 20 g/L NaCl (Fig. 1d, ANOVA, F_3,8_ = 137.1, *p* < 0.001). The requirement for NaCl on N_2_ fixation was further verified by the direct quantification of N_2_ fixation activities. In accordance with the growth pattern (Fig. 1d), N_2_ fixation activities were almost completely inhibited at low NaCl concentrations (0, 2 g/L) and maximised at 18 g/L NaCl (approximately 75 nmol ethylene produced per 10^8^ cells per hour) (Fig. 1e, ANOVA, F_3,8_ = 318.3, *p* < 0.001). These results are of particular interest, as they contrast sharply with the conclusions of previous studies that coastal N_2_ fixation is sensitive to NaCl^2,3^.

**Fig. 1.**
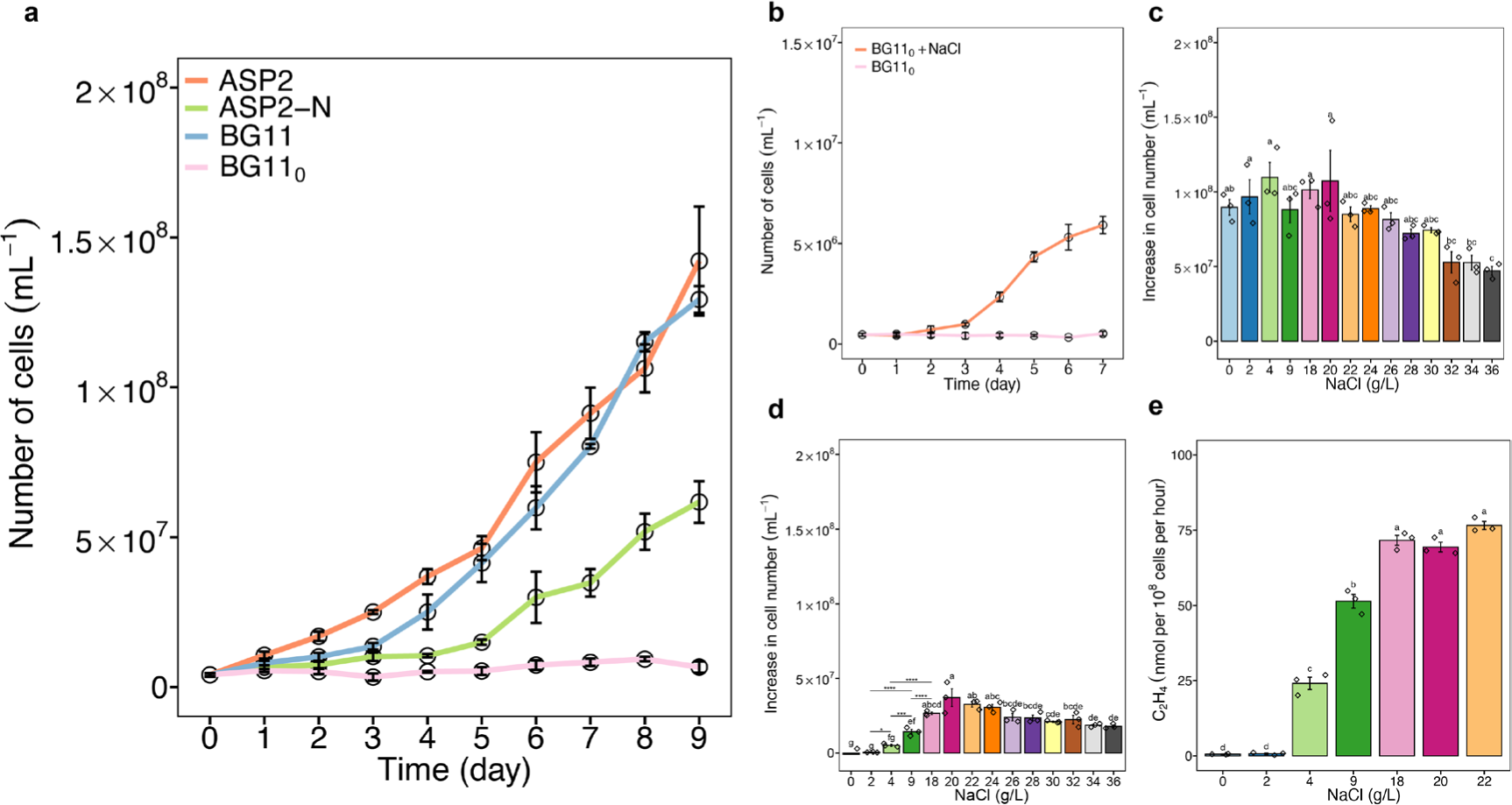
Growth response to NaCl under N-replete or N-depleted conditions. **a** Growth curves of cells grown in four different media over a period of 9 days. **b** Growth curves of seven-day-old non-growing cells in BG11_0_ with the addition of NaCl over a period of 7 days. **c** Cell number increase of cells grown in BG11 with NaCl gradients. **d** Cell number increase of cells grown in BG11_0_ with NaCl gradients. **e** N_2_ fixation activity of cells grown in BG11_0_ with NaCl gradients. Nitrogenase activity is expressed as ethylene (C_2_H_4_) production by acetylene reduction. Data are presented as mean ± standard deviation of triplicate values. Different letters on each bar (**c**, **d**, **e**) indicate statistical significance (*p* < 0.05) calculated by one-way ANOVA with Tukey’s HSD post-hoc analysis over all populations tested. In figure **d**, line segments and corresponding significance markers indicate statistical significance calculated by one-way ANOVA across four salinity treatments (2, 4, 9, 18 g/L). Significance: ns (no significance), *(*p* < 0.05), **(*p* < 0.01), ***(*p* < 0.001), ****(*p* < 0.0001).

### Cellular N starvation due to lack of N_2_ fixation

To test whether these non-growing cells in BG11_0_ are physiologically N-starved due to the non-functional N_2_ fixation, nitrogenous compounds and dry weight of cells cultured in either BG11_0_ or BG11_0_ (NaCl) were quantified. Consistent with our expectation, compared to normally growing cells (functioning N_2_ fixation) with NaCl, NaCl deprivation led to a significant decrease in chlorophyll (C_55_H_72_O_5_N_4_Mg) (*t* test, *p* < 0.05), total protein content (*t* test, *p* < 0.01), dry weight (*t* test, *p* < 0.01), and total protein ratio (of dry weight) (*t* test, *p* < 0.05) of non-N_2_-fixing cells grown in BG11_0_ (Extended Data Fig.1a-d).

### Cellular N-starvation causes a stringent response

Transcriptomic analysis was performed to reveal the molecular background and transcriptomic features of the NaCl requirement for N_2_ fixation. Compared to N_2_-fixing cells cultured with NaCl (18 g/L), 5226 (1909 genes upregulated, 3317 genes downregulated) genes of non-N_2_-fixing cells grown without NaCl were identified as differentially expressed genes (DEGs) (adjusted *P* value < 0.05) (Extended Data Fig. 2a). According to KEGG enrichment analysis, all DEGs could be classified into several functional categories (Extended Data Fig. 2b), showing striking metabolic differences. Detailed heatmap visualisation of DEGs enriched in KEGG ribosome biosynthesis (map03010, Fig. 2a), porphyrin metabolism (participating in chlorophyll synthesis) (map00860, Extended Data Fig. 2c), and photosynthesis (map00195, Extended Data Fig. 2d) showed that almost all DEGs enriched of non-N_2_-fixing cells showed a decreased pattern at the transcript level compared to N_2_-fixing cells. Given the importance of ribosomes and photosynthesis for cell growth and division, these distinct gene expression patterns provide clues at the molecular level as to why diazotrophic cells cultured without NaCl fail to grow or divide.

**Fig. 2.**
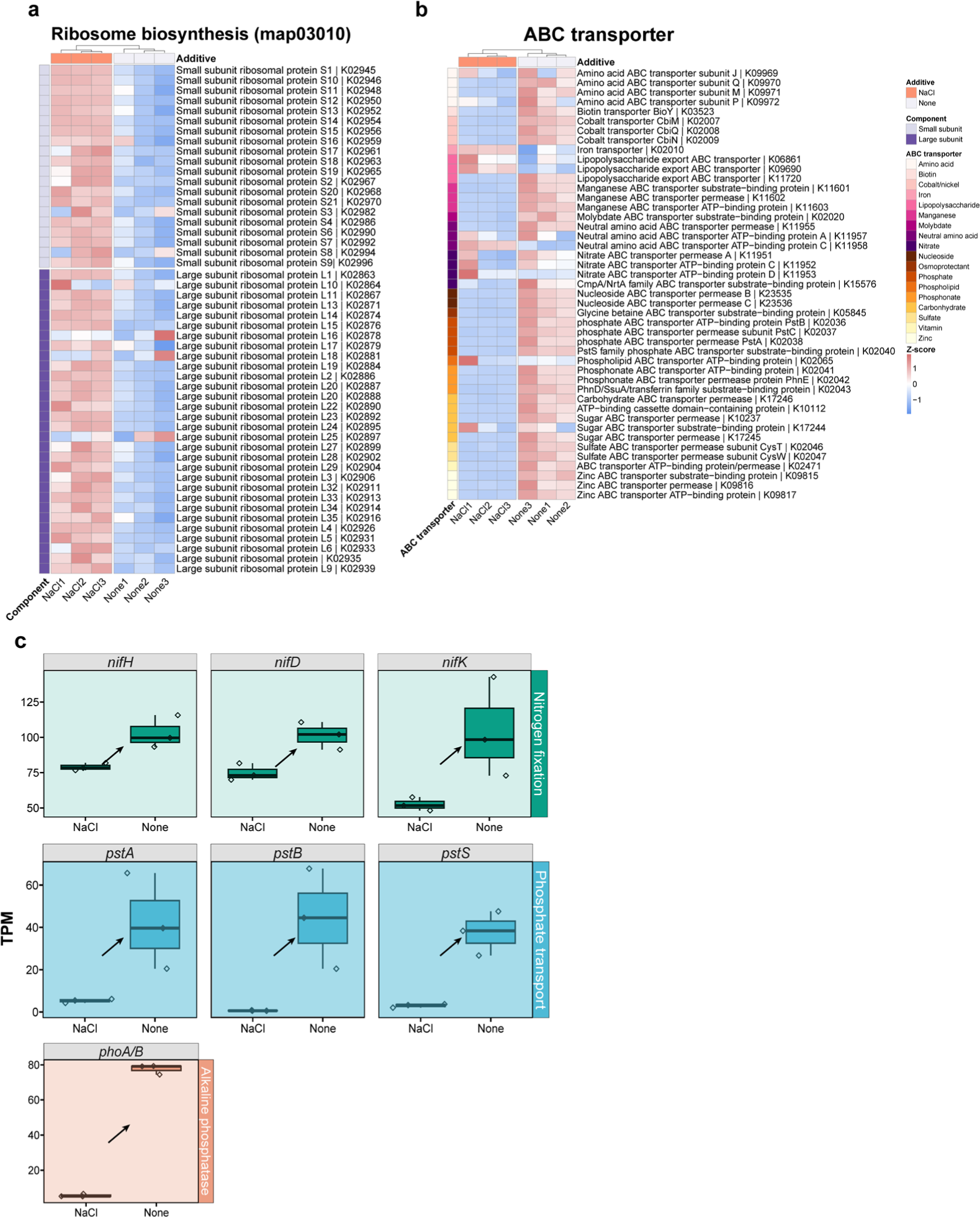
Enrichment analysis of ribosome biosynthesis and nutrient transporter genes, and gene expression trend of nitrogenase, P limitation biomarkers. **a** Heatmap analysis showing the DEGs enriched for ribosome biosynthesis and **b** for nutrient transporters. Two components of the ribosomal proteins (small subunit, large subunit) were analysed, and eighteen transporter types were annotated and compared. Column dendrograms indicate similarity based on Euclidean distance and hierarchical clustering. Gene clusters were determined by k-means clustering with Euclidean distance. “NaCl” denotes the addition of extra NaCl (18 g/L) to BG11_0_, while “None” indicates that no additional substances were added to BG11_0_. The heatmap colour gradient shows low gene expression (blue) and high gene expression (red). **c** Expression trend of nitrogenase structural genes (*nifHDK*) and P limitation biomarkers (*pstA, pstB, pstS, phoA/B*). Shaded-in plots indicate significantly different expression levels of genes and arrows indicate the directionality of statistically significant trends between these two treatments (*p* < 0.05). Statistical significance of selected genes was calculated from TPM (transcripts per million) pairwise comparisons of triplicate samples of NaCl deprivation treatment (None) compared to the NaCl addition treatment (NaCl).

It is reasonable to expect that non-N_2_-fixing cells (N-starved) cells will reduce their primary metabolism to a minimal level in order to survive. Indeed, these cells exhibited a stringent response^10^ (a persistent state, in which cells reduce protein synthesis and stop dividing), as evidenced by their cellular N starvation (Extended Data Fig. 1), repression of ribosome biosynthesis (Fig. 2a), upregulated expression of *relA* (encoding ppGpp, signalling molecule for the stringent response^10^) (Extended Data Fig. 3), upregulated nutrient transporter genes, a transcriptional indicator of the stringent response^11^ (Fig. 2b, 32 out of 45 annotated genes), and growth arrest (Fig. 1a, d).

Regarding the nitrogenase (key enzyme responsible for N_2_ fixation) structural genes (*nifHDK*)^12^, to our surprise, we found even higher expression of *nifHDK* in the non-N_2_-fixing cells, i.e. cultured without NaCl, than in the N_2_-fixing cells, cultured with NaCl (Fig. 2c). As P is another essential nutrient for living cells, we also focused on the transcript pattern of commonly used P limitation biomarkers, the phosphate binding transporter genes (*pstA, pstB, pstS*) and the alkaline phosphatase gene (*phoA/B*)^13^. Consistent with the stringent response theory, higher expressions of all four P limitation biomarkers were observed in these non-N_2_-fixing cells (Fig. 2c). This upregulated pattern of these genes makes biological sense: in a stringent response, starving cells struggle to meet nutrient demands by expressing higher levels of nutrient synthases or transporters. However, it should be noted that the stringent response described above is the consequence (resulted from cellular N starvation) rather than the cause of N_2_ fixation dysfunction in the absence of NaCl; how N_2_-fixers are N-starved in the absence of NaCl remains to be determined.

### N_2_-fixation depends exclusively on Na^+^

To determine whether the need for NaCl is specific, we tested whether four other metal chlorides, including KCl, LiCl, MgCl_2_ and CaCl_2_, at doses equivalent to NaCl, could stimulate the growth of cells grown in BG11_0_ in the same way as NaCl. Since N_2_ fixation is a must for population growth under N deprivation, population growth is used as a proxy for N_2_ fixation activity in the following contexts, unless otherwise specified. We found that population growth was only detected in the NaCl treatment, indicating that the requirement for NaCl is exclusive and cannot be substituted by other metal chlorides (Fig. 3a, ANOVA, F_4,10_ = 65.74, *p* < 0.001). Moreover, since all the tested metal chlorides contain a chloride moiety, this result also indicates that Na^+^ and not Cl^−^ plays a role in N_2_ fixation. Notably, BG11_0_ medium alone contains 0.082 g/L Na^+^ from its constitutive ingredients (Na_2_CO_3_, Na_2_EDTA·2H_2_O and Na_2_MoO_4_·2H_2_O), nevertheless, this Na^+^ concentration is far below the effective concentration for activating N_2_ fixation as shown earlier (Fig. 1d, e).

**Fig. 3.**
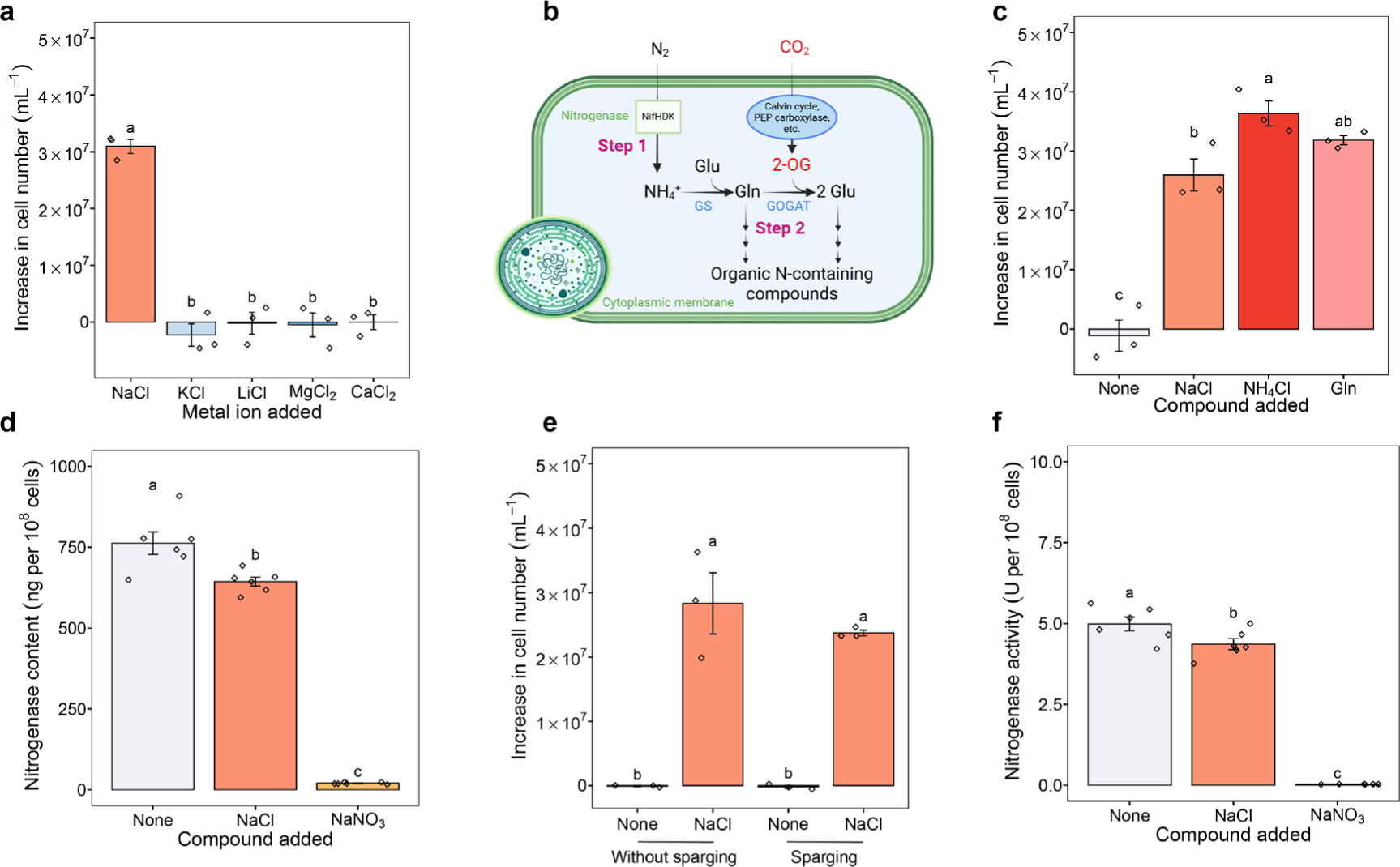
Possible mechanisms underlying the NaCl requirement for N_2_ fixation. **a** Increase in cell number of cells grown in BG11_0_ with the addition of five different metal chlorides: NaCl, KCl, LiCl, MgCl_2_ and CaCl_2_. **b** Schematic representation of the pathway of unicellular cyanobacterial N_2_ fixation. N_2_ is first reduced to NH_4_^+^ by nitrogenase (Step 1) and then is incorporated into carbon skeletons via the glutamine (Gln) synthetase-glutamate (Glu) synthase pathway, where further N-containing organic metabolites are formed (Step 2). NifHDK, nitrogenase complex; PEP carboxylase, phosphoenolpyruvate carboxylase; 2-OG, 2-oxoglutarate; GS, glutamine synthetase; GOGAT, glutamate synthase. **c** Cell number increase of cells grown in BG11_0_ amended with NaCl, NH_4_Cl and Gln. **d** Nitrogenase content of cells grown in BG11_0_, BG11_0_ (NaCl) and BG11_0_ (NaNO_3_). **e** Cell number increase of cells grown in BG11_0_ and BG11_0_ (NaCl) with and without N_2_ sparging. **f** Nitrogenase activity of cells grown in BG11_0_, BG11_0_ (NaCl) and BG11_0_ (NaNO_3_). One unit (U) is the amount of enzyme that catalyses the reaction of 1 µmol of substrate per minute. The graphs show the mean ± standard deviation, n = 3 for **a**, **c** and **e**, n = 6 for **d** and **f**. Different letters on each bar represent statistical significance (*p* < 0.05) calculated by one-way ANOVA with Tukey’s HSD post-hoc analysis over all populations tested.

### Conformationally active nitrogenase despite lack of Na^+^

Unicellular cyanobacterial N_2_ fixation can generally be divided into two steps, i.e. N_2_ fixation (from gaseous N_2_ to ammonium) and N assimilation (from ammonium to organic nitrogenous compounds)^14^ (Fig. 3b). We have shown that N_2_ fixation is severely inhibited (Fig. 1e), but it remains to be determined whether the synthesis of organic N-containing metabolites is also suppressed under NaCl deprivation. Therefore, we performed a nutrient addition experiment in which the product of N_2_ fixation (ammonium (NH_4_Cl)) and an intermediate of N assimilation (glutamine (Gln)) was added to cells grown in BG11_0_. Compared to cells grown in BG11_0_, the addition of NH_4_Cl and Gln mimicked the Na^+^ effect and resulted in normal growth (Fig. 3c, ANOVA, F_3,8_ = 58.83, *p* < 0.001), confirming that N_2_ fixation only was inhibited by Na^+^ deprivation.

We then tested: 1)whether Na^+^ deficiency inhibited the biosynthesis of nitrogenase; 2) whether Na^+^ plays a role in protecting the enzyme from oxygen (O_2_) inactivation, as nitrogenase is known to be extremely sensitive to O_2_^15^; 3) whether Na^+^ participates in the conformational activation of nitrogenase. Although almost no N_2_ fixation activity was detected in cells grown in BG11_0_ (Fig. 1e), these cells expressed a significantly higher amount of nitrogenase (Fig. 3d, ANOVA, F_2,15_ = 337.6, *p* < 0.001), which is consistent with our previous transcriptome analysis showing that non-N_2_-fixing cells expressed higher levels of *nifHDK* (Fig. 2c). The results of the sparging experiment show that even under anaerobic conditions, Na^+^ was still required for N_2_ fixation and population growth (Fig. 3e, ANOVA, F_3,8_ = 40.52, *p* < 0.001), suggesting that the dependence of nitrogenase activity on Na^+^ is not due to a role of the cation in providing oxygen protection to the enzyme. Nitrogenase from non-N_2_-fixing cells was tested to be conformationally active with a significantly higher catalytic rate compared to that of N_2_-fixing cells (Fig. 3f, ANOVA, F_2,15_ = 289.2, *p* < 0.001), which makes sense since the catalytic capacity of an enzyme is generally proportional to its amount (Fig. 3d). Taken together, these results indicate that: 1)Na^+^ is not involved in the biosynthesis, protection against O_2_, inactivation or conformational activation of nitrogenase; 2) non-N_2_-fixing cells grown without Na^+^ synthesised an unusually high amount of nitrogenase.

### A shortage of ATP prevents N_2_ fixation

Biological N_2_ fixation, including both N_2_ fixation itself (the theoretical minimum energy requirement) and the associated apparatuses supporting N_2_ fixation, is highly energy-intensive^16^, and we therefore reasoned that a limitation of ATP caused by Na^+^ deficiency might lead to N_2_ fixation dysfunction. To test this hypothesis, we grew cells in BG11_0_ supplemented with extra ATP. The results show that extra ATP stimulated population growth (Fig. 4a, ANOVA, F_2,6_ = 49.95, *p* < 0.001), suggesting that: 1) the lack of ATP could be the underlying mechanism why the nitrogenase is not working; 2) Na^+^ is involved in ATP biosynthesis under diazotrophic conditions.

**Fig. 4.**
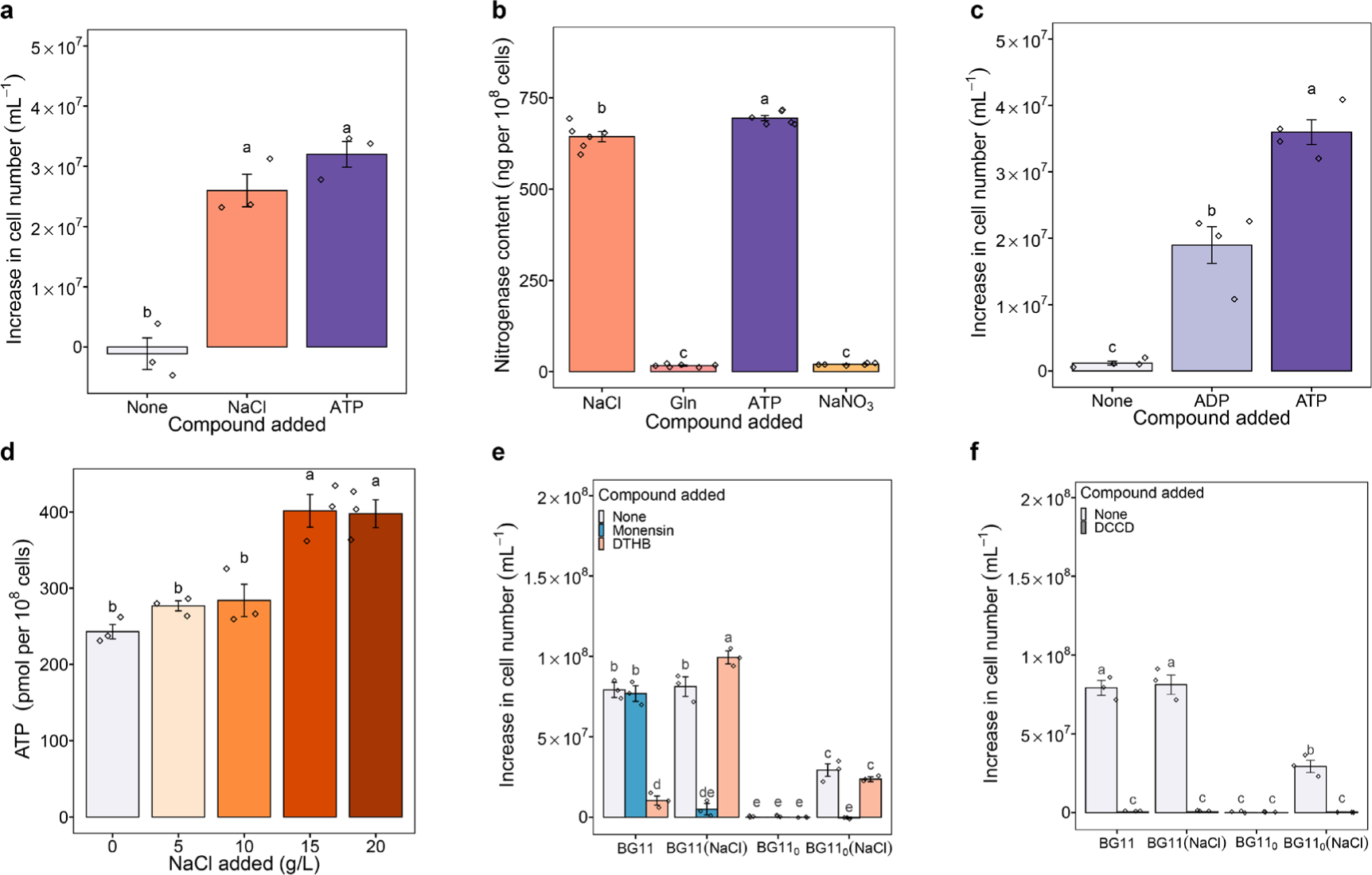
N_2_ fixation & population growth with ATP supplementation, cellular ATP quantification, monensin, DTHB and DCCD tests. **a** Increase in cell number of cells grown in BG11_0_ or supplemented with either NaCl or ATP. **b** Nitrogenase content of cells grown in BG11_0_ (NaCl), BG11_0_ (Gln), BG11_0_ (ATP) and BG11_0_ (NaNO_3_). **c** Increase in cell number of cells grown in BG11_0_ supplemented with equivalent doses of ATP or ADP. **d** Cellular ATP content of N-starved cells after the addition of NaCl. **e** Increase in cell number of cells grown in the medium supplemented with monensin and DTHB as compared to non-supplemented cells. **f** Cell number increase of cells grown in medium supplemented with DCCD as compared to non-supplemented ones. The graphs show the mean ± standard deviation, n = 3 for **a**, **d, e** and **f**, n = 4 for **c**, and n = 6 for **b**. Different letters on each bar represent statistical significance (*p* < 0.05) calculated by ANOVA with Tukey’s HSD post-hoc analysis across all populations tested.

However, the presence of the nitrogen element in ATP (C_10_H_16_N_5_O_13_P_3_) makes us reconsider the role of ATP in activating N_2_ fixation, either by providing chemical energy or by providing N. To distinguish between these two possibilities, we first examined the nitrogenase content of cells grown in BG11_0_ supplemented with extra ATP. Almost no nitrogenase was observed in cells supplemented with N (Gln, NaNO_3_), whereas cells supplemented with ATP and Na^+^ synthesised the two highest levels of nitrogenase (Fig. 4b, ANOVA, F_3,20_ = 2269, *p* < 0.001). It thus can be concluded that ATP here functions as available energy other than providing N. In parallel, we also assessed population growth in BG11_0_ supplemented with an equivalent dose of ATP or ADP (adenosine diphosphate). As shown in Fig. 4c, cells with extra ATP had the significantly highest growth in all treatments (ANOVA, F_2,9_ = 81.24, *p* < 0.001), indicating that ATP acts as extra chemical energy rather than as an N resource, which would otherwise result in similar growth as both additives provide an equivalent dose of N. From an energetic point of view, this result makes sense, as ATP carries more energy than ADP, and can therefore provide more power for the nitrogenase to fix more N_2_ for growth. All results indicate that insufficient bioavailable ATP (energy) leads to N_2_ fixation dysfunction in the absence of Na^+^. However, how Na^+^ is involved in the energy metabolism of N_2_-fixing cells remains unclear.

### Na^+^ energetics enables N_2_ fixation

Regarding energy metabolism in bacteria, it is well established that ATP synthesis is driven by the energy stored in a transmembrane electrochemical gradient of protons coupled to the light-driven or the respiration-driven proton pumps (H^+^ energetics)^17^. Nevertheless, there is evidence that Na^+^ can replace H^+^ as the coupling ion to drive ATP synthesis in several anaerobic bacteria^17,18^. This type of machinery for synthesising ATP by Na^+^-coupling (Na^+^ energetics) fits well with our findings that Na^+^ correlates with the supply of ATP for N_2_ fixation. In particular, there is a striking resemblance between N_2_-fixing *Cyanothece* sp. and anaerobic bacteria with Na^+^ energetics, as both require non-oxic microenvironments. Therefore, we propose the existence of a mechanism for Na^+^-coupled ATP synthesis in N_2_-fixing *Cyanothece* sp.. To test this proposal, we quantified the cellular ATP content of N-starved cells grown in BG11_0_ (seven days) ten minutes after the addition of NaCl. Higher concentrations of NaCl (15 and 20 g/L) resulted in a significantly higher amount of ATP relative to no and low NaCl addition (5 and 10 g/L) (Fig. 4d, ANOVA, F_4,10_ = 19.93, *p* < 0.001), directly confirming the role of Na^+^ in driving ATP synthesis.

To better understand the role of Na^+^ energetics in energy metabolism, we investigated cell growth after adding either a Na^+^-specific ionophore monensin^19,20^, a protonophore DTHB^21^, or a typical F-type ATP synthase inhibitor DCCD^22^ to the respective culture medium. If Na^+^ fluxes were coupled to ATP synthesis, the collapse of the Na^+^ transmembrane gradient caused by monensin should lead to growth arrest due to the inhibition of ATP synthesis. Compared to normally growing cells from the control, 14 µM monensin significantly inhibited the growth of cells in BG11_0_ (NaCl) (Fig. 4e, ANOVA, F_2,24_ = 190.6, *p* < 0.001), whereas cells cultured in BG11 with monensin divided normally. Only these cells grown in BG11 were significantly inhibited by DTHB (200 µM). No growth was observed in all treatments treated with DCCD (100 µM) (Fig. 4f, two-way ANOVA, F_1,16_ = 1420.4, *p* < 0.001). Together, these results indicate: 1) the existence of a Na^+^-coupled ATP synthesis machinery, as monensin inhibited the growth of cells in BG11_0_ (NaCl); 2) the existence of an H^+^-coupled ATP synthesis machinery, as cells grew normally in BG11 unaffected by monensin, but were severely impaired by DTHB; and 3) that Na^+^ and H^+^ energetics use the same type of F_0_F_1_ ATP synthase, as DCCD suppressed the growth of cells in all treatments. Interestingly, we found that cells in BG11 (NaCl) with monensin were unable to grow. This suggests that Na^+^ availability may play a critical role in determining the coupling ion for ATP synthase. In this case, Na^+^-coupled ATP synthesis would be favoured by the presence of Na^+^, otherwise, the cells in BG11 (NaCl) with monensin should have grown due to H^+^ energetics. Overall, these results suggest an essential role for Na^+^ energetics in activating N_2_ fixation.

### Na^+^ gradient generators

Similar to the H^+^ cycle, a Na^+^ cycle is required for continuous energy supply^17,18^. In this case, Na^+^ pumps (Na^+^ gradient generators) must work together with ATP synthase (Na^+^ gradient consumer) to re-energise the membrane in N_2_-fixing cells. Two potential Na^+^ pumps were identified by transcriptomic analysis: the Na^+^-motive ferredoxin:NAD oxidoreductase (Rnf)^23^, originally identified for its role in N_2_-fixing *Rhodobacter* ^24^, and the Na^+^/H^+^ antiporter. We found that in the absence of Na^+^, non-N_2_-fixing cells expressed significantly higher levels of genes encoding these two pumps (Extended Data Fig. 4). This suggests that these cells cannot establish a Na^+^ gradient to drive ATP synthesis to meet their energy needs.

### Sodium energetics in cyanobacteria

Sodium energetics was initially thought to occur only in some extremophilic and anaerobic bacteria^18,25–27^. This paradigm is changing with the realisation that this type of membrane bioenergetics is more than an adaptation to extreme environments, but probably the most ancient form, originating in the Last Universal Cellular Ancestor (LUCA)^28^. As the first membranes were permeable to protons, the ion gradient was based on the impermeability of the membrane to sodium^18,29^. Accordingly, it has recently been shown that sodium-based bioenergetics is widespread throughout the tree of life (based on genome sampling for Na^+^-specific ATPases^30^). Nevertheless, Na^+^ energetics has not been much discussed in the cyanobacterial phyla, but see its role in cyanobacterial dormancy^20^. To investigate the prevalence of Na^+^ energetics in cyanobacteria, sequence alignment analysis was performed with a focus on the ion binding sites of the c ring (AtpH)^20,31,32^ of the ATP synthase, where small amino acid sequence variations around the ion binding site determine whether it binds H^+^ or Na^+^ ions^20,33^.

Here, we show a summary of the H^+^ selectivity results based on sequence alignment of *Cyanothece* sp. AtpH protein sequences with these of *Ilyobacter tartaricus* (low H^+^ selectivity) with published crystal structures^31^ and 111 other cyanobacterial species (Fig. 5a provides a summary of the data, but details of the alignment can be checked in Extended Data Fig. 5). H^+^ selectivity is defined in three subcategories: Low+Medium/High (all species with low H^+^ selectivity harbour two AtpH homologs - one with low (Na^+^ energetics) and the other with medium or high H^+^ selectivity (both types of energetics)), Medium (medium H^+^ selectivity, capable of both types of energetics) and High (high H^+^ selectivity, i.e. capable of H^+^ energetics only). Interestingly, we find that 78 out of 112 species are capable of Na^+^ energetics (Extended Data Fig. 5). This indicates their energetic plasticity with coupling ions, as evidenced by their medium or low H^+^ selectivity, i.e., the coupling ion of their ATP synthase can be either H^+^ or Na^+^. As expected, in freshwater species, none of the 63 species analysed showed low H^+^ selectivity, while 45 species showed medium H^+^ selectivity (capable of both H^+^ and Na^+^ energetics) and 18 species showed high H^+^ selectivity (capable of H^+^ energetics only). In marine/brackish species, although cells typically encounter low H^+^ and high Na^+^ environments, 16 out of 50 species still showed high H^+^ selectivity (capable of H^+^ energetics only).

**Fig. 5.**
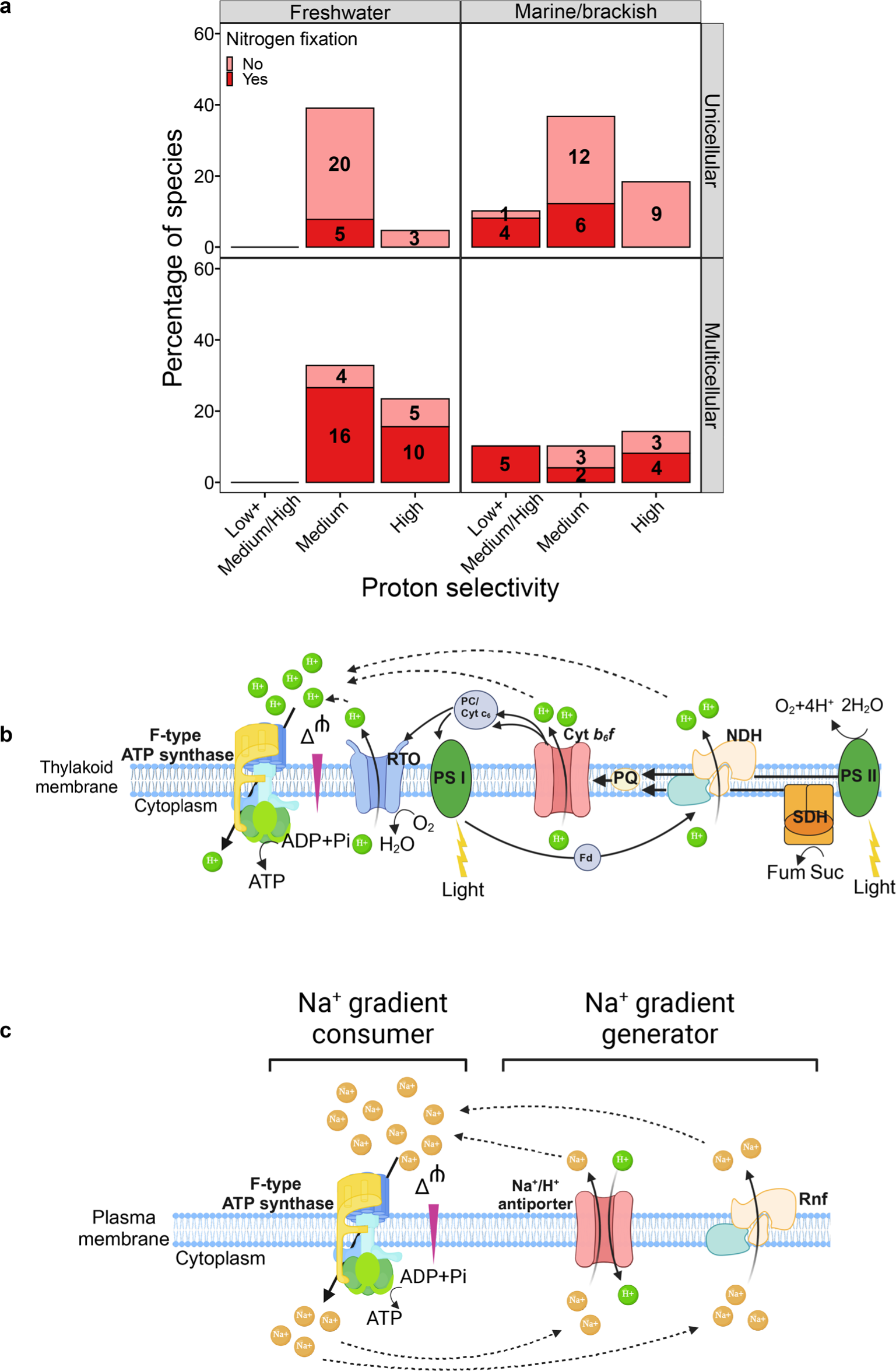
H^+^ and Na^+^ energetics in cyanobacteria. **a** A summary of the proton selectivity of AtpH from 112 cyanobacterial species based on AtpH sequence alignment. The percentage of species (y-axis) indicates the ratio of species with different proton selectivity to habitat-based categories (freshwater, 63 species; marine, 49 species). Low+Medium/High (all species with low proton selectivity harbour two AtpH homologs - one with low and the other with medium or high H^+^ selectivity), Medium (medium H^+^ selectivity) and High (high H^+^ selectivity). The numbers on the graph represent the number of species with the corresponding proton selectivity. **b** Schematic representation of the typical H^+^ energetics across the thylakoid membrane coupled to photosynthesis in cyanobacteria (including *Cyanothece* sp.). **c** Schematic representation of the proposed Na^+^ energetics across the plasma membrane with Na^+^ gradient generator and consumer in N_2_-fixing *Cyanothece* sp. cells. Abbreviations: RTO, respiratory terminal oxidase; Cyt *b_6_f*, cytochrome *b_6_f*; PC/Cyt c_6_, plastocyanin/cytochrome c_6_; PQ, plastoquinone; Fd, flavodiiron; NDH, NADH-dehydrogenase; Fum, fumarate; Suc, succinate; SDH, succinate dehydrogenase; Glu, glutamic aid; Gln, glutamine; Ser, Serine; Val, valine; Tyr, tyrosine; Ala, alanine; ATP, Adenosine triphosphate; ADP, Adenosine diphosphate; Pi, inorganic phosphate; Rnf, Na^+^-translocating ferredoxin:NAD oxidoreductase; Δψ, membrane potential.

All 15 unicellular species capable of N_2_ fixation, including 10 marine/brackish species and 5 freshwater species, are capable of Na^+^ energetics. This is likely due to the energy challenges faced in nature, particularly under diazotrophic conditions, and highlights the critical role of Na^+^ energetics in N_2_ fixation. While none of the unicellular N_2_-fixing species rely solely on H^+^ energetics (high H^+^ selectivity), 10 out of 26 freshwater and 4 out of 11 marine/brackish filamentous N_2_-fixing species exclusively use H^+^ energetics. This finding makes sense as energy generated by photosynthesis of filamentous species in the light can directly power N_2_ fixation^15,34^.

Interestingly, we found 10 marine/brackish species (9 N_2_-fixing species, one of which is *Cyanothece* sp.), that possess two AtpH homologs. One homolog has low H^+^ selectivity, while the other has medium/high H^+^ selectivity, suggesting that these species can conduct both Na^+^ and H^+^ energetics.

Based on the evidence provided, we suggest that *Cyanothece* sp. carries out typical H^+^ energetics across the thylakoid membrane (Fig. 5b). During the night under N_2_-fixing conditions, however, we propose that the primary Na^+^ gradient consumer (Na^+^-coupled ATP synthase) works together with active Na^+^ gradient generators (Rnf and Na^+^/H^+^ antiporter) across the plasma membrane to drive ATP synthesis for N_2_ fixation (Fig. 5c).

### Anaerobic respiration rather than aerobic respiration possibly powers N_2_ fixation

To understand why *Cyanothece* sp. cells capable of H^+^ energetics must perform Na^+^ energetics under N_2_-fixing conditions, it is important to note that N_2_ fixation in this species occurs in the dark and under anaerobic conditions in the cell^16^. Although H^+^ energetics is generally believed to be more energetically advantageous than Na^+^ energetics, under these conditions, H^+^ energetics is no better than Na^+^ energetics due to the lack of high potential electron acceptor, i.e., O_2_^18^. Therefore we speculate that unicellular N_2_ fixation is driven by fermentation, rather than aerobic respiration as previously postulated^15,34^. Although our transcriptomic analysis was designed to distinguish non-N_2_-fixing growth-arrested cells from normally growing N_2_-fixing cells at the transcript level, we also identified interesting transcript patterns of fermentation-related genes that indicate that fermentation, rather than aerobic respiration, generates energy for N_2_ fixation. Specifically, we found that three key genes involved in microbial fermentation^35^, namely *ack* encoding acetate kinase, *adh* encoding alcohol dehydrogenase and *ldh* encoding lactate dehydrogenase, were upregulated in non-N_2_-fixing cells compared to N_2_-fixing cells (Extended Data Fig. 6a). This suggests that non-N_2_-fixing cells may face challenges in meeting their energy requirements in the absence of Na^+^ and therefore express higher levels of key fermentative energy metabolism genes. The presence of fermentative energy generation in N_2_-fixing cells is further supported by the significantly higher enzyme activities in N_2_-fixing than that in cells grown with N (Extended Data Fig. 6b).

### High salinities do not coincide with high temperatures

In nature, N_2_ fixation and growth of coastal N_2_ fixers are influenced by several environmental factors, singly or in combination. In particular, temperature affects the development of cyanobacterial populations^36,37^, which also indirectly affects salinity dynamics. For cyanobacteria, higher temperatures (> 25°C, with optimal growth temperature at 30°C for *Cyanothece* sp.)^36,38^ are preferred for growth. We have already shown that N_2_ fixation of *Cyanothece* sp. depends on higher salinity, but in nature, N_2_-fixation-favouring higher salinity and growth-favouring higher temperature usually don’t coincide, because higher temperature seasons are usually accompanied by higher precipitation, resulting in higher coastal freshwater input (fluviatile and terrestrial freshwater) and thus a lower salinity^39,40^. We analysed the salinity dynamics of eleven stations along the Texas Gulf coast (Extended Data Fig. 7a), where *Cyanothece* sp. was originally isolated, over the past decade. The annual surface seawater temperatures of the selected stations were also processed and compared. As expected, although varying between stations, the annual salinity dynamics at most sites follow the trend that higher salinities occur during months with lower temperatures, while lower salinities occur during months with higher temperatures (Extended Data Fig. 7b,c). Thus, higher temperature months are often linked with higher precipitation, leading to decreased salinities and lower N_2_ fixation and growth *in situ*. In controlled experimental setups, low temperatures (20°C and 10°C) severely inhibited population development. Even when N was abundant, populations under low-temperature regimes grew less than diazotrophic populations at 30°C (Extended Data Fig. 7d, ANOVA, F_1,12_ = 82.37, *p* < 0.001), demonstrating the critical role of temperature on N_2_ fixation and growth. This may further explain the low abundance of cyanobacterial N_2_-fixers in N-deficient coastal waters.

## Discussion

In coastal aquatic ecosystems, N is often a limiting nutrient^2,3^. It would therefore be logical to expect N_2_-fixing cyanobacteria to thrive under these conditions. However, a puzzling ecological paradox emerges as N_2_-fixing cyanobacteria, despite their ecological advantage in N-deficient environments, maintain a low abundance in coastal waters^1–3^. This study challenges the traditional view that coastal N_2_ fixation is sensitive to high levels of NaCl^2,3^. Instead, we report that N_2_ fixation by a coastal unicellular N_2_-fixer is exclusively dependent on the presence of Na^+^. In the absence of Na^+^, the cells failed to fix N_2_ and showed a stringent response, despite producing substantially higher levels of conformationally active nitrogenase. Further experiments show that the presence of Na^+^ is a prerequisite for the activation of Na^+^ energetics for the generation of ATP, which is subsequently used for the energy-intensive N_2_ fixation. Thus, this study demonstrates the exclusive and essential role of Na^+^ energetics in powering cyanobacterial N_2_ fixation in likely all uni- and some multicellular species.

Although three species of filamentous N_2_-fixing cyanobacteria were reported to be dependent on NaCl for N_2_ fixation about forty years ago^41^, it was concluded at that time that inhibition of P uptake in the absence of Na^+^ was responsible for the loss of nitrogenase activity. However, this cannot explain the Na^+^ dependence of N_2_ fixation in *Cyanothece* sp., as we show here that P uptake is independent of the presence of Na^+^ (Fig. 1a, Fig. 3c). Only recently, although not directly related to N_2_ fixation, a study using *Synechocystis* sp. PCC 6803 reported that N-starved cells engage in Na^+^ energetics for ATP synthesis for maintaining viability and to awaken from dormancy^20^. These reports suggest that Na^+^ energetics may be more widespread in cyanobacteria than commonly thought. This is supported by our finding of the high prevalence of Na^+^ energetics capabilities in cyanobacteria based on the AtpH sequence alignment analysis (Fig. 5 and Extended Data Fig. 5). According to our analysis, 78 out of 112 species are capable of Na^+^ energetics, and all 15 unicellular N_2_ fixers exhibit this ability. In fact, Na^+^ energetics is considered to be the most ancient form of bioenergetics. Genomic analyses suggest that it is still prevalent throughout the tree of life today^18,30^.

Conventionally, H^+^ energetics are thought to be chemically more advantageous than Na^+^ energetics^18^. So why should cells capable of H^+^ energetics perform Na^+^ energetics? In *Cyanothece* sp., as in all unicellular N_2_-fixing cyanobacteria and in some multicellular species, energy-intensive N_2_ fixation occurs in the dark^16^. The necessary ATP is believed to be produced through H^+^ energetics, coupled with the respiratory breakdown of endogenous storage carbohydrates. It is thought that aerobic respiration of the stored glycogen granules is critical for N_2_ fixation at the beginning of the dark period, not only because it energises this process but also because it consumes intracellular O_2_, creating an anoxic interior (so-called respiratory protection)^15,34^. While an anoxic environment is required for a functional nitrogenase to fix N_2_, this process is very energy-intensive. Thus, if this energy were produced solely by aerobic respiration of the glycogen granules, cells would need to increase respiration and hence their respiratory O_2_ uptake. Simultaneously, the O_2_ level in the cell should be kept low enough to avoid inactivation of nitrogenase. Consequently, the demand for more O_2_ for increased aerobic respiratory activity is in conflict with the demand for an anaerobic interior. These two processes, which should work in tandem, are incompatible. Therefore, it is reasonable to assume that under N_2_ fixation conditions (dark and anoxic), aerobic respiration fuelled by H^+^ energetics is not favoured and instead Na^+^ energetics, which can function independently of aerobic respiration^17^, appears to be an alternative.

Furthermore, under dark and anaerobic conditions, Na^+^ energetics are not inferior to H^+^ energetics from an energetic point of view. Only when there is a direct mechanistic link between redox reactions (oxidation of external resources) or photosynthesis in photosynthetic microbes to the translocation of H^+^ across the membrane, H^+^ energetics is chemically more advantageous than Na^+^ energetics^18^. Thus, an energetic advantage only exists when high potential electron acceptors, such as O_2_, are available. This is probably why H^+^ is mainly used as the coupling ion in the aerobic microbial world. We propose that for *Cyanothece* sp., the presence of O_2_ determines which type of energetics is used. Under conditions where N is abundant, O_2_ does not interfere with cellular metabolism during the day or night, allowing for the more efficient use of H^+^ energetics. However, when N is depleted, an obligate anaerobic environment is required for oxygen-sensitive N_2_ fixation. In this case, Na^+^ energetics is used as O_2_, the electron acceptor for proton energetics, is unavailable.

Functioning Na^+^ energetics requires both a Na^+^ gradient consumer (ATP synthase) and Na^+^ gradient generators (Na^+^ pumps). We have identified two Na^+^ pumps, the Na^+^/H^+^ antiporter and Rnf (Extended Data Fig. 4), in N_2_-fixing *Cyanothece* sp.. However, which type of energy metabolism drives the Na^+^ gradient generators when aerobic respiration is excluded? Anaerobic respiration, i.e. fermentation, is the possible choice. Indeed, under dark and anaerobic environments (identical to the conditions required for unicellular N_2_ fixation), it has been shown that several cyanobacteria use fermentation for energy generation and survival^35^. Although our focal organism, *Cyanothece* sp., is known to possess all the genes required for fermentation^42^, fermentation has never been demonstrated or postulated to occur in the context of N_2_ fixation. Nevertheless, we suggest here that fermentation, as reported in other anaerobic N_2_-fixing microbes^43,44^, likely powers Na^+^ pumps and ultimately N_2_ fixation in unicellular phototrophic cyanobacteria such as *Cyanothece* sp.. This is supported by the upregulated expression of unique fermentation-related genes and significantly higher activities of corresponding enzymes (Extended Data Fig. 6).

Interestingly, most of the filamentous cyanobacteria also do not rely on aerobic respiration in the dark for N_2_ fixation, but gain the energy from photosystem I-mediated cyclic phosphorylation in the light^16^. This is true not only for heterocystous cyanobacteria, but also for the non-heterocystous *Trichodesmium* sp., where aerobic respiration in the light is normally inhibited, and the unfavoured O_2_ is consumed via the Mehler reaction^45^.

From an energy perspective, the advantage of engaging both H^+^ and Na^+^ energetics in *Cyanothece* sp. can be rationalised by considering the energy challenges faced by cells in their complex natural habitats. To meet the energy demands of dynamic coastal seas, particularly the energy-intensive anaerobic N_2_ fixation, organisms require a versatile energy metabolism. i.e., H^+^ energetics coupled to photosynthesis and aerobic respiration during the day and Na^+^ energetics for the anaerobic N_2_ fixation in the dark. This metabolic flexibility likely contributes to the evolutionary hardiness of cyanobacteria in a wide range of environments.

Contrary to the conventionally accepted paradigm that high salinity inhibits coastal N_2_ fixation, this study reports that N_2_ fixation exclusively depends on Na^+^. This provides insights into how salinity modulates coastal N_2_ fixation and partially explains why N_2_-fixing cyanobacteria cannot thrive in N-limited coastal waters. Moreover, the fact that low NaCl levels (caused by freshwater input) during periods of growth-supporting high temperatures, which are unable to support N_2_ fixation, may contribute to the low abundance of N_2_-fixing unicellular cyanobacteria in coastal seas. However, in the context of global climate change, current patterns of change (global warming and drought^6,46^) may result in periods optimal for both N_2_ fixation and population growth, likely catalysing the global expansion of cyanobacterial blooms. Furthermore, the reported Na^+^ energetics provide experimental evidence for their existence in microbial phototrophs in general, and the first evidence for their role in cyanobacterial N_2_ fixation in particular. This provides another way of understanding the energy metabolism of N_2_ fixation in unicellular cyanobacteria. However, it also suggests that sodium energetics may still be more widespread than expected since its probable evolutionary origin in LUCA and that its metabolic importance in other organisms may still be underestimated.

## Materials and methods

### Strain and culture conditions

*Cyanothece* sp. ATCC 51142 (recently reclassified as *Crocosphaera subtropica*^47^) was obtained from the ATCC culture collection. Stock cells were grown photoautotrophically at continuous light with a light intensity of 30 μmol m^−2^ s^−1^ in artificial seawater medium ASP2^9^ at 30°C. All experiments were carried out in a light-dark cycle of 12 h/12 h to accommodate the temporal segregation of photosynthesis and N_2_ fixation, unless stated otherwise.

### N-dependent effects of NaCl on population growth

To evaluate the effect of NaCl on the growth and the N_2_ fixation activity of *Cyanothece* sp., we performed growth experiments in which we grew cells in artificial seawater medium ASP2 and artificial freshwater medium BG11^48^, supplemented with or without N, respectively. In total, we had four media, i.e. ASP2, ASP2 without N (ASP2-N), BG11 and BG11 without N (BG11_0_). Prior to conducting the growth experiments, exponentially growing stock cells were harvested by centrifugation, washed twice to remove NaCl and resuspended in these four sterile media. The resuspended cells were subsequently inoculated into tissue culture flasks (Thermo Scientific, USA) containing 10 mL of the respective medium with a starting density of approximately 4*10^6^ cells/mL. Cultures were grown aerobically at a light intensity of 30 μmol m^−2^ s^−1^ in a light-dark cycle of 12 h/12 h on a shaker at 150 rpm (stroke: 26 mm, MQD-S3R, Shanghai Minquan Instruments Co., Ltd) at 30°C (unless otherwise specified, the same culture parameters were used in all experiments). Growth experiments lasted for nine days, and we recorded cell density daily using a cell counting chamber (Neubauer haemocytometry) under a light microscope (Axio observer Z1, Carl Zeiss, Germany).

To investigate the effects of NaCl deficiency on growth in the absence of N, cells previously grown in BG11_0_ for seven days were subcultured in BG11_0_ supplemented with either NaCl (18 g/L) or NaNO_3_ (1.5 g/L or 17.6 mM). BG11_0_ without extra additives was used as a blank control. Each culture was inoculated with a starting density of ∼ 5*10^5^ cells/mL. Growth experiments were conducted in triplicate over seven days. Cell density was quantified as described above.

To further investigate the effect of NaCl on the growth of *Cyanothece* sp. in the presence of N, we conducted experiments using three replicates (with an initial density of approximately 4*10^6^ cells/mL) for each of the fourteen NaCl concentrations (0, 2, 4, 9, 18, 20, 22, 24, 26, 28, 30, 32, 34, 36 g/L) in BG11 medium. The experiments lasted for seven days, after which cell density was quantified and the increase in cell number was determined. Similar experiments were conducted using BG11_0_ medium instead of BG11 to evaluate the impact of NaCl on the growth of *Cyanothece* sp. in the absence of N. These experiments lasted for seven days, during which cell density and population growth were recorded and calculated. The population growth (the increase in cell number) was determined by subtracting cell density on day 0 from the cell density on day 7.

### Measurement of nitrogenase activity

Nitrogenase activity of cells cultured in BG11_0_ with seven NaCl concentrations (0, 2, 4, 9, 18, 20, 22 g/L) was measured by a modified acetylene reduction method. The cultures were grown for five days under the conditions described above, and the cell density was quantified on the last day. The assays were conducted in Vacutainer tubes containing 2 mL of the corresponding cell samples and 15% acetylene in the gas phase. Each sample was incubated for 6 hours under the same conditions of light, temperature, and shaking as those of the growth experiments. 0.2 mL of triplicate samples were analysed for their ethylene concentration by high-performance liquid chromatography (HPLC). The rates of nitrogenase activity are reported in nanomoles of C_2_H_4_ produced per 10^8^ cells per hour.

### Cellular chlorophyll, total protein, dry weight

To quantify cellular chlorophyll, 1 mL of cells of *Cyanothece* sp. cell cultures aged 7 days grown in either BG11_0_ or BG11_0_ with 18 g/L of NaCl medium (hereafter BG11_0_ (NaCl)), was centrifuged and then extracted twice with 5 mL of 80% aqueous acetone. The extracts were pooled, and the spectra of this extract and a sample of whole cells were measured using a DW2000 spectrophotometer (Olis, GA, USA), both against 80% acetone or BG11 media as a reference. Chlorophyll *a* and *b* levels from the acetone extracts were calculated as outlined in a previous study^49^. Specifically, the concentration of chlorophyll was determined using the equations: Chl *a* (µg/mL) = 12.25A_663_ –2.79A_647_ and Chl *b* (µg/mL) = 21.5A_647_ – 5.1A_663_ (A_n_ is the absorbance spectrophotometrically measured at the specific wavelength of 663 and 647). These values were used to calculate total chlorophyll = Chl *a* + Chl *b*. All measurements were done in triplicate.

Total protein content was determined using a Bradford Protein Assay kit (Solarbio, China) following the manufacturer’s instructions. Cells grown in either BG11_0_ or BG11_0_ (NaCl) for 7 days were collected and cell density was recorded. Then, 2 mL of culture from each treatment was used to extract total protein using a Plant Protein Extraction Kit (Solarbio, China). Specifically, 2 mL of sample was centrifuged and the supernatant was discarded. Glass beads were used to grind samples and break the cell wall thoroughly under liquid nitrogen. 1 mL of lysis solution was added at 4°C for 20 minutes, samples were shaken every 5 minutes, followed by centrifugation at 4°C at 20000 × g for 30 minutes. The resulting supernatant was harvested for quantification. All experiments were done in triplicate.

To quantify cell dry weight, 10 mL of cells grown in either BG11_0_ or BG11_0_ (NaCl) were concentrated in pre-weighed Eppendorf tubes and cell density was quantified. Then, each treatment was centrifuged at 20000 × g for 5 minutes. The resulting pellet was washed twice with ddH_2_O and the supernatant medium was discarded. The tubes were dried in an oven at 65°C for 24 hours, and the total dry weight (cells + tubes) was subsequently determined gravimetrically. Cellular dry weight was calculated with the following equation: Cellular dry weight = (Total dry weight – Weight of tube) / cell number.

### Metal chlorides, NH_4_Cl, Glutamine and ATP addition

To explore the mechanisms of growth inhibition in the absence of NaCl under diazotrophy, we grew cells in BG11_0_ and provided them with additional substances to determine which part of the metabolism is affected by NaCl deficiency. To this end, we assessed whether NaCl could be replaced by other metal chlorides by adding an equivalent amount (310 mM) of KCl, LiCl, MgCl_2_ and CaCl_2_. To investigate whether N_2_ fixation products can lead to population growth, we supplemented the BG11_0_ medium with additional NH_4_Cl (2 mM) or glutamine (2 mM, Aladdin, China), respectively. We conducted 7-day-long growth experiments using three replicates (with an initial density of approximately 4*10^6^ cells/mL) with these experimental setups, cell densities were recorded and the increases in cell number were calculated.

### Sparging experiment

For anaerobic incubation, cells with a starting density of approximately 4*10^6^ cells/mL were transferred into tissue culture flasks with either BG11_0_ or BG11_0_ supplemented with 18 g/L NaCl. In total, four treatments were included, i.e. sparging (BG11_0_, BG11_0_ (NaCl)) and no sparging (BG11_0_, BG11_0_ (NaCl)). For the two sparging treatments, tissue culture flasks were flushed with N_2_ for 10 minutes and then sealed with parafilm to prevent diffusive influx of O_2_. Triplicate cultures were cultivated for seven days under the conditions above. Afterwards, the cell density of each treatment was quantified.

### Quantification of nitrogenase content and estimation of nitrogenase activity

The enzyme content and activities of nitrogenase of *Cyanothece* sp. collected in the dark period were measured using commercial ELISA (enzyme-linked immunosorbent assay) kits (Shanghai Ruifan Biological Technology Co., Ltd, Shanghai, China) with corresponding model substrates (six replicates per treatment) according to the manufacturer’s instructions. The NaNO_3_ treatment is used to assess the accuracy of the ELISA approach, as cells should not synthesise nitrogenase when N is replete. Enzymes were extracted from cells grown in the respective media (approximately 1*10^8^ cells) in an Eppendorf tube containing 1 mL of ice-cold PBS buffer (pH = 7.4). The homogenate was centrifuged at 1200 × g for 20 minutes at 4 °C and the resulting supernatant was used directly for spectrophotometric assays.

### Transcriptomics

20 mL of cells grown in either BG11_0_ or BG11_0_ with NaCl (18 g/L) were harvested in the dark phase and then centrifuged at 20000 × g for 10 minutes at 4 °C to remove the supernatant. Cell pellets were resuspended in ddH_2_O and centrifuged again. The supernatants were discarded to remove residual medium. This procedure was repeated twice. The collected cell pellets were transferred to RNase-free cryogenic vials and fast frozen in liquid nitrogen for 30 minutes until extraction. In total, we had both cell types in triplicate.

Total RNA was extracted using a Plant RNA Purification Reagent for plant tissue (Invitrogen, USA), and genomic DNA was removed using DNase I (TaKara, Dalian). Only high quality RNA samples (OD260/280 = 1.8∼2.2; OD260/230 ≥ 2.0; RIN ≥ 6.5; 28S:18S ≥ 1.0; and > 1μg) were used for sequencing library construction. Total RNA samples were sent to Shanghai Majorbio Bio-pharm Technology Co., Ltd. (Shanghai, China). Paired-end sequencing was performed on the Illumina sequencing platform (San Diego, CA, USA) according to the manufacturer’s instructions (Illumina).

Data were analysed using the online platform Majorbio Cloud (www.majorbio.com)^50^. Raw paired-end reads were trimmed and quality controlled for adapter contamination using SeqPrep (https://github.com/jstjohn/SeqPrep) and Sickle (https://github.com/najoshi/sickle) with default parameters. Clean reads were then aligned to the *Cyanothece* sp. ATCC 51142 genome using HISAT2 software (http://ccb.jhu.edu/software/hisat2/index.shtml). Mapped reads from each sample were assembled using StringTie (https://ccb.jhu.edu/software/stringtie/index.shtml?%20t=example) in a reference-based approach.

To identify the differentially expressed genes (DEGs) between the two different treatments, the expression level of each transcript was calculated using the transcripts per million reads (TPM) method. RSEM (http://deweylab.biostat.wisc.edu/rsem/) was used to quantify gene abundance. Essentially, differential expression analysis was performed using DESeq2 with a Q-value ≤ 0.05, and significant DEGs were confirmed only if |log2FC| > 1 and a Q-value ≤ 0.05. Finally, KEGG pathway analyses were performed using Goatools (https://github.com/tanghaibao/Goatools) and KOBAS (http://kobas.cbi.pku.edu.cn/home.do), respectively. TPM was used for data visualisation. Heatmaps were generated using ComplexHeatmap v2.8.0. Heatmap Z-scores were calculated for each gene by subtracting the gene expression from the row mean and then dividing by the row standard deviation. All other plots were generated using ggplot2 v3.3.5.

### ATP addition experiment

To examine whether the inhibition of nitrogenase activity was caused by the lack of ATP, additional ATP (0.2 mM, Solarbo, China) was added to BG11_0_. We tested the growth of three replicates (with an initial density of approximately 4*10^6^ cells/mL) in the modified BG11_0_ media. After seven days, cell density was quantified and the increase in cell number was calculated.

To determine the actual role of ATP in activating population growth, cells (approximately 4*10^6^ cells/mL) were cultured in BG11_0_ modified with NaCl (18 g/L), glutamine (2 mM), ATP (0.2 mM), and in BG11, respectively. Each treatment had six replicates. Cells were harvested in the dark period on day 7 for quantification of nitrogenase content by ELISA according to the manufacturer’s protocol (Shanghai Ruifan Biological Technology Co., Ltd, Shanghai, China). If the role of ATP is to provide chemical energy, then nitrogenase biosynthesis would be expected; if ATP acts as an N resource, then nitrogenase synthesis is unnecessary.

In addition, an equivalent dose (0.2 mM) of ADP (Macklin, China) and ATP was added to the cells grown in BG11_0_. Each treatment had four replicates. After seven days, the increase in cell number was calculated and contrasted. If the extra ATP acts as combined N, population growth would be expected to be similar between the ATP and ADP treatments, as these two treatments contain equivalent amounts of N. On the contrary, since ATP contains higher energy than ADP^51^, higher population growth from the ATP treatment will provide evidence that ATP acts as the energy supply.

### ATP quantification

To test whether Na^+^ directly mediates the intracellular ATP content, we quantified the cellular ATP content of cells in different NaCl gradients. Cells with a starting density of 4*10^6^ cells/mL were first cultured in BG11_0_ for seven days. Since N_2_ fixation occurs in the dark, NaCl was added to these non-growing cultures in the dark phase on day 7. Five NaCl concentrations (0, 5, 10, 15, 20 g/L) were included. 10 minutes after the addition of NaCl, cells were fast frozen in liquid nitrogen for 20 minutes and stored at −80°C until ATP quantification by ELISA (MEIMIAN, Jiangsu Meimian Industrial Co., Ltd, China) according to the manufacturer’s protocol.

### Monensin, DTHB, and DCCD test

The following assays were performed to understand the role of Na^+^ or H^+^-coupled ATP synthesis in driving ATP generation in the presence or absence of Na^+^ under diazotrophy. The Na^+^-specific ionophore monensin (Solarbo, China) and the protonophore 3,5-di-tert-butyl4-hydroxybenzaldehyde (DTHB, Thermo Fisher Scientific, USA) were used to test which pathway drives ATP synthesis. In addition, a typical F_0_F_1_ ATP synthase inhibitor, dicyclohexylcarbodiimide (DCCD, Macklin, China), was used to test whether the H^+^-motive and Na^+^-motive machinery share a common ATP synthase. We had four media, including BG11, BG11 with NaCl (18 g/L), BG11_0_, BG11_0_ with NaCl (18 g/L), and cells (with a starting density of approximately 4*10^6^ cells/mL) were cultured in each treatment with either monensin (14 µM), DTHB (200 µM), DCCD (100 µM) or no additive (as blank control). In total we had twelve treatments with three replicates each. After seven days of growth, the increases in cell number were recorded and contrasted.

### AtpH sequence alignment

In a typical F_0_F_1_-type ATP synthase, binding ion specificity is determined by the c ring (AtpH) of the F_0_ complex^31,32^. Both H^+^ and Na^+^ bind a glutamate residue, and small differences in the amino acid sequence around the ion binding site determine the preference of the c-ring, for either H^+^ or Na^+^. For example, the neighbouring groups of Na^+^ ATP synthase are generally polar, whereas H^+^ ATP synthase favours hydrophobic groups^20,31^. The balance between polar and hydrophobic residues makes the c-rings more or less selective for one ion or the other. In the model organism *I. tartaricus* with strong Na^+^ selectivity but low H^+^ selectivity, in addition to the main ion-bind glutamic acid (E 65), 5 key neighbouring amino acids (3 Na^+^-binding residues: Q 32, V 63, S 66; 2 water-binding residues: A 64, T 67 determine the high Na^+^ selectivity (low H^+^ selectivity) of the strain’ AtpH.

We included 112 cyanobacterial species, isolated from typical freshwater (63 species) and marine/brackish habitats (49 species), with available AtpH sequence. FASTA type of data of AtpH from all the bacterial strains analysed was downloaded from UniProtKB (https://www.uniprot.org/help/uniprotkb). Sequence alignment analysis was performed using Jalview software (https://www.jalview.org/). The structure image of the c-ring of AtpH from *I. tartaricus* was generated on the Protein Data Bank website (https://www.rcsb.org/).

### Annual salinity and temperature analysis

Annual salinity records for the last ten years from eleven selected stations along the Gulf of Mexico coasts were downloaded from the Coastal Salinity Index website (https://www2.usgs.gov/water/southatlantic/projects/coastalsalinity/index.html), and annual surface seawater records were downloaded from the US National Centers for Environmental Information (https://www.ncei.noaa.gov/access/coastal-water-temperature-guide/all_table.html#egof). The stations used are: Calcasieu River (Cameron); Vermilion Bay (Cypremort Point); Caillou Bay (Cocodrie); Hackberry Bay (Grand Isle); Barataria Bay (Grand Isle); Barataria Bay (Grand Terre Island); Black Bay (Snake Island); Mississippi Sound (Joseph Island); Mississippi Sound (Grand Pass); Mississippi Sound (Merrill Shell); Biloxi Bay (Cadet Harbour).

### Population growth at different temperatures

To determine the role of temperature on the population development of *Cyanothece* sp., 10 mL of cells (a starting density of approximately 4*10^6^ cells/mL) were grown in either BG11 with NaCl (18 g/L) or BG11_0_ with NaCl (18 g/L) under three temperature regimes: 10°C, 20°C and 30°C. This growth experiment was run for seven days, and the increase in cell number was calculated and contrasted.

### Estimation of the activity of fermentation-related enzymes

To determine the activities of fermentation-related enzymes, cells (approximately 4*10^6^ cells/mL) were cultured in BG11 with NaCl (18 g/L) and in BG11_0_ with NaCl (18 g/L), respectively. Each treatment had nine replicates. The experiment was carried out in a 12h/12h light-dark cycle, and cells were harvested twice on day 7 at midnight to estimate the activities of three fermentation-related enzymes using ELISA, according to the manufacturer’s protocol (Jiangsu Meimian Industrial Co., Ltd, Yancheng, China). Three enzymes were acetate kinase, alcohol dehydrogenase, and lactate dehydrogenase.

### Statistical analysis

Physiological parameters were analysed using Welch’s *t* test. Growth data, nitrogenase activity data (the acetylene reduction method) and ELISA data were analysed by one-way ANOVA. Monensin and DTHB data were analysed by three-factor ANOVA. DCCD data was analysed by two-way ANOVA. Significance: ns (no significance), *(*p* < 0.05), **(*p* < 0.01), ***(*p* < 0.001), ****(*p* < 0.0001). All statistical analyses were performed in R v4.1.1^52^.

## Acknowledgements

We would like to thank Yuelu Jiang for her helpful comments on this study. This work was supported by the S&T Projects of Shenzhen Science and Technology Innovation Committee (JCYJ20200109142822787, KCXFZ2022, RCJC20200714114433069, and JCYJ20200109142818589), the Project of Shenzhen Municipal Bureau of Planning and Natural Resources (Grant No. [2021]735-927), as well as the Shenzhen-Hong Kong-Macao Joint S&T Project (pending number 202205303000176). K.H. gratefully acknowledges the financial support of the John Templeton Foundation (#62220). The opinions expressed in this paper are those of the authors and not those of the John Templeton Foundation.

## Author contributions

S.T., K.H. and ZH.C. conceptualised and designed the experiments. S.T., XY.C., YQ.L., L.L., Q.Y., D.L., JM.Z. (Jianming Zhu) performed the experiments. S.T., K.H. and J.Z. (Jin Zhou) analysed and visualised the data. All authors interpreted the results and wrote the manuscript.

## Data availability

All data obtained for visualisation and statistics in this study are accessible online at Zenodo (https://doi.org/10.5281/zenodo.10046202). Raw RNAseq reads for the analysis of differential gene expression have been submitted to NCBI’s SRA database (http://www.ncbi.nlm.nih.gov) under BioProject PRJNA1014498. Source data are provided with this paper.

## Code availability

All scripts for data visualisation are available at https://github.com/SiTANG1990/N2-fixation-coastal-unicellular-cyanobacteria.git.

