## Supplementary Information for "Unicellular cyanobacteria rely on sodium energetics to fix N_2_"

[1] Shenzhen Public Platform for Screening and Application of Marine Microbial Resources, Tsinghua Shenzhen International Graduate School, Shenzhen, Guangdong Province, PR China.

[2] Institute of General Microbiology, Kiel University, Kiel, Germany.

[3] Technology Innovation Center for Marine Ecology and Human Factor Assessment of Natural Resources Ministry, Tsinghua Shenzhen International Graduate School, Shenzhen, 518055, Guangdong Province, PR China.

* Correspondence:
 (K.H.), (ZH.C.)

Includes extended data figures 1-7

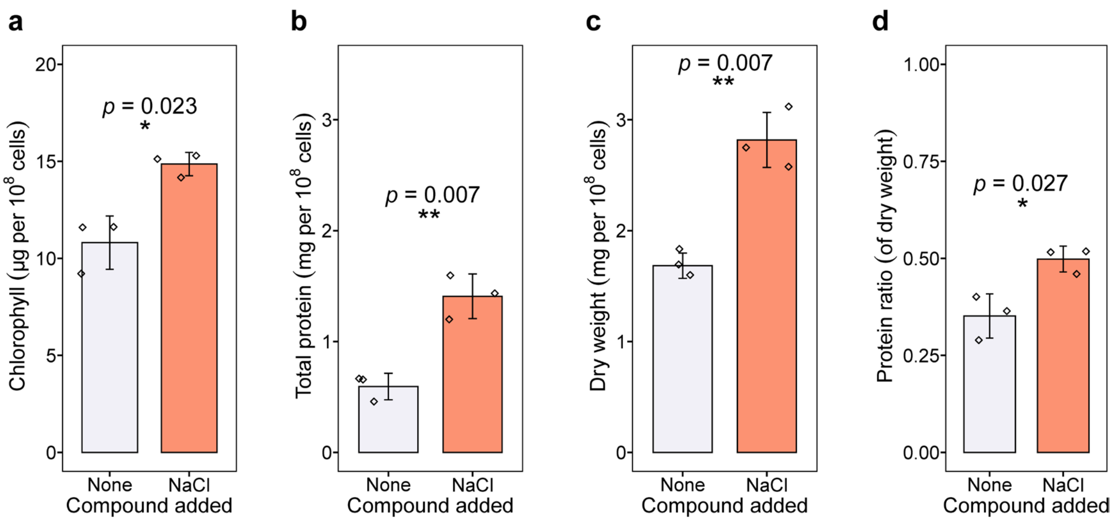


**Extended Data Fig. 1. Physiological parameters in the absence or presence of NaCl under diazotrophy. a** Chlorophyll content. **b** Total protein. **c** Dry weight. **d** Protein ratio of dry weight. “None” indicates that no additional substances were added to $BG11_0$ while “NaCl” denotes the addition of extra NaCl (18 g/L) to $BG11_0$. The graphs show the mean ± standard deviation of the triplicate values. Corresponding significance markers represent statistical significance calculated by Welch’s *t* test. Significance: ns (no significance), *($p$ < 0.05), **($p$ < 0.01), ***($p$ < 0.001), ****($p$ < 0.0001).

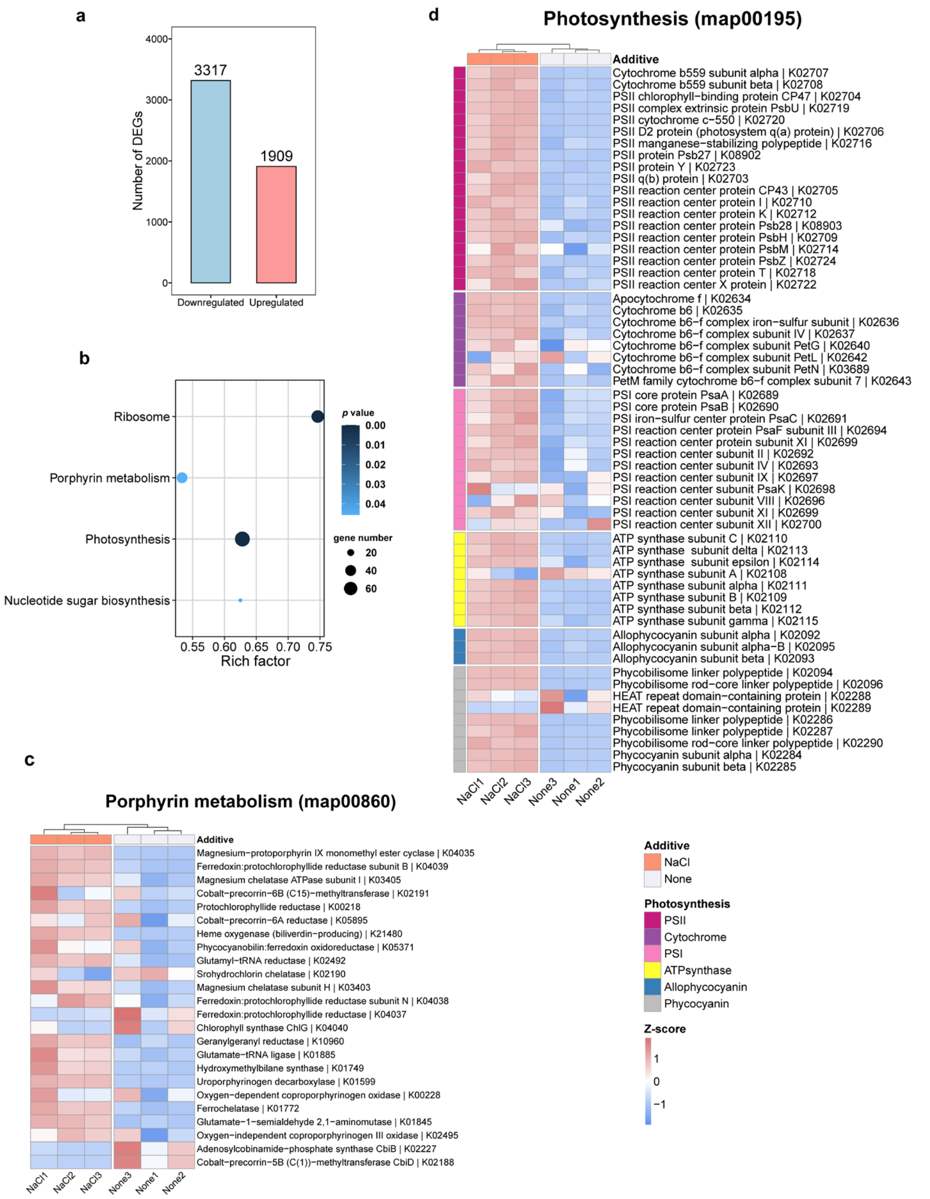


**Extended Data Fig. 2. Transcriptomes and key functional enrichment assay of DEGs. a** Number of significantly upregulated or downregulated DEGs (number of genes denoted at the top). **b** Analysis of enriched KEGG Pathways containing DEGs with a *p* value < 0.05. The size of the dot reflects the number of genes for each pathway, and the colour reflects the *p* value. **c** Heatmap analysis showing the DEGs enriched for porphyrin metabolism (map00860) and **d** for photosynthesis (map00195). Six photosynthetic subunits (photosystem II

(PSII), cytochrome, photosystem I (PSI), ATP synthase, allophycocyanin, phycocyanin) were analysed. Column dendrograms show similarity based on Euclidean distance and hierarchical clustering. Gene clusters were determined by k-means clustering with Euclidean distance. “NaCl” denotes the addition of NaCl (18 g/L) to $BG11_0$ while “None” indicates that no additional substances were added to $BG11_0$. KEGG annotations were assigned from the genome annotation. The heatmap colour gradient shows low gene expression (blue) and high gene expression (red).

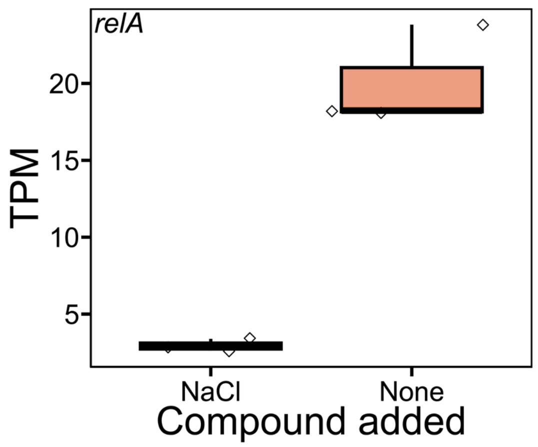


**Extended Data Fig. 3. Transcript analysis of the stringent response signal-encoding gene *relA*.** Expression trend of the stringent response signal ppGpp encoding gene *relA*. *relA* was identified as a significantly different expression gene between these two treatments ($p$ < 0.05). Statistical significance of *relA* was calculated from TPM (transcripts per million) pairwise comparisons of triplicate samples of NaCl deprivation treatment (None) compared to the NaCl addition (18 g/L) treatment (NaCl).

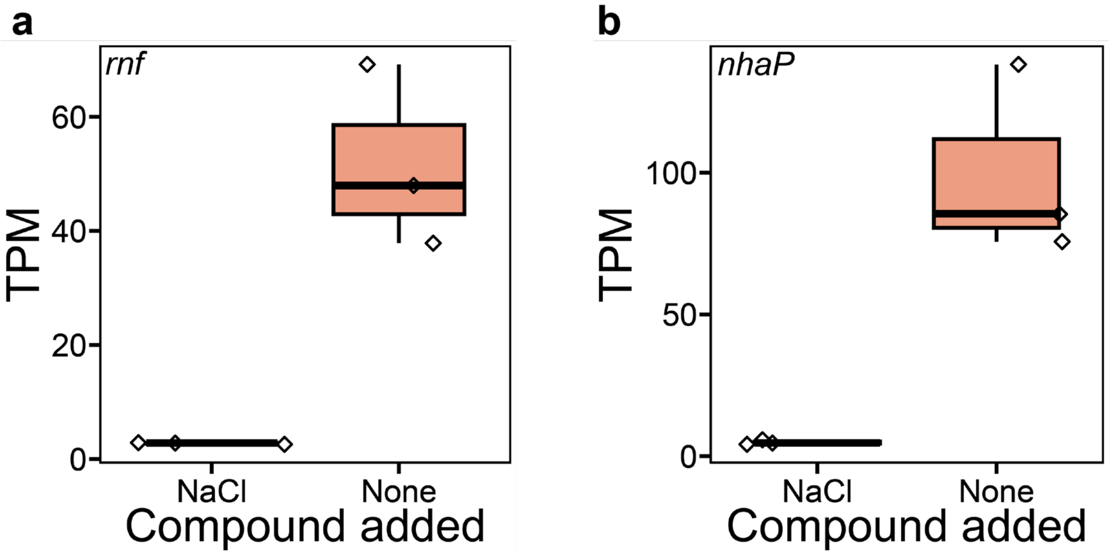


**Extended Data Fig. 4. Expression trend of the $Na^+$ pump Rnf encoding gene *rnf* (a) and $Na^+/H^+$ antiporter encoding gene *nhaP* (b).** Both genes were identified as significantly different expression genes between these two treatments ($p$ < 0.05). Statistical significance of *rnf* and *nhaP* was calculated from TPM (transcripts per million) pairwise comparisons of triplicate samples of NaCl deprivation treatment (None) compared to the NaCl addition (18 g/L) treatment (NaCl).

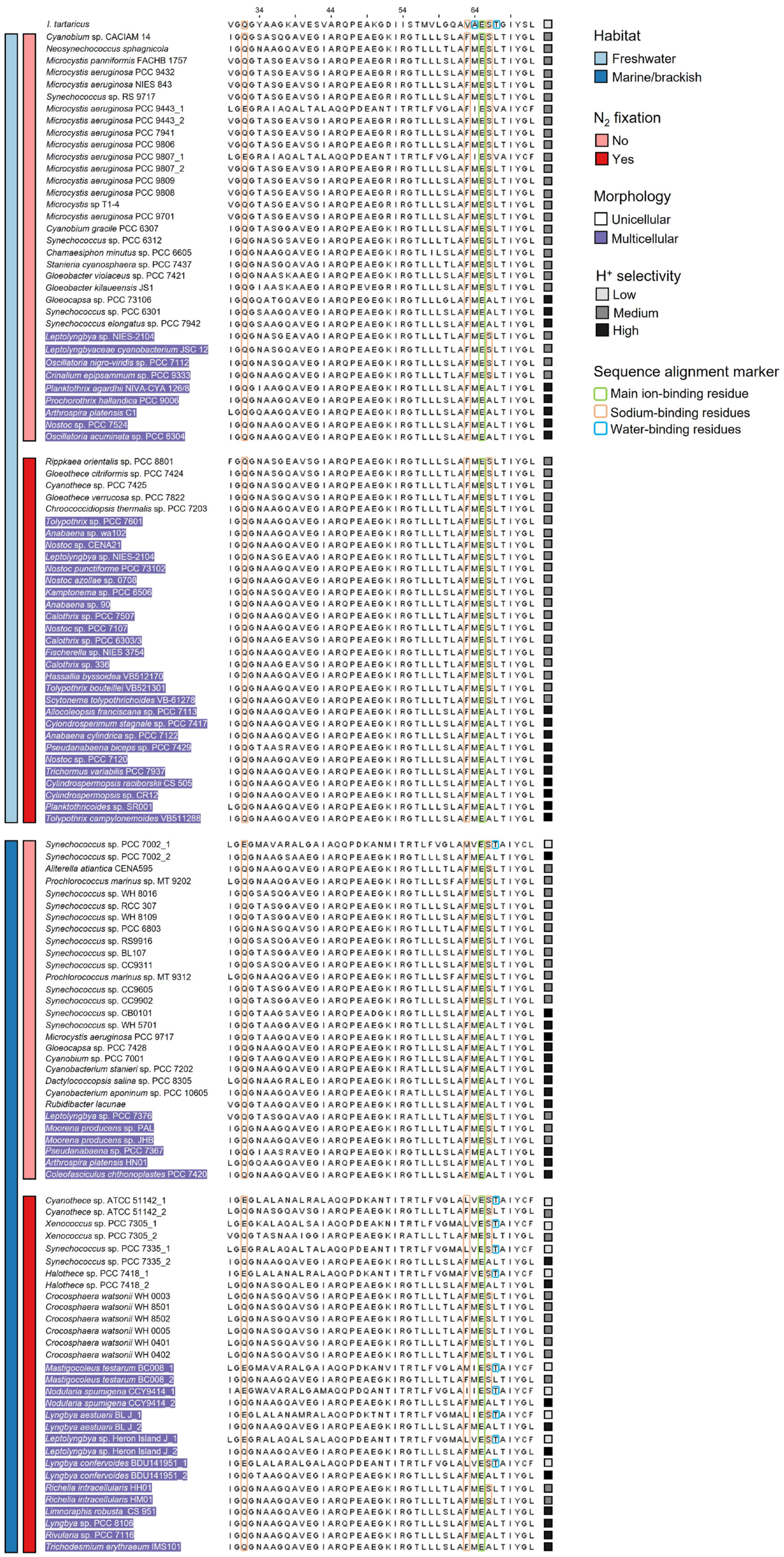
34
44
54
64
Habitat
Freshwater
Marine/brackish
$N_2$ fixation
No
Yes
Morphology
Unicellular
Multicellular
$H^+$ selectivity
Low
Medium
High
Sequence alignment marker
Main ion-binding residue
Sodium-binding residues
Water-binding residues

**Extended Data Fig. 5. Sequence alignment of the ion-binding site of AtpH from *Ilyobacter tartaricus* with 112 cyanobacterial species.** Amino acid residues involved in $Na^+$ ion coordination, habitat, $N_2$ fixation ability, morphology and proton selectivity of the analysed cyanobacterial species are marked with different colours.

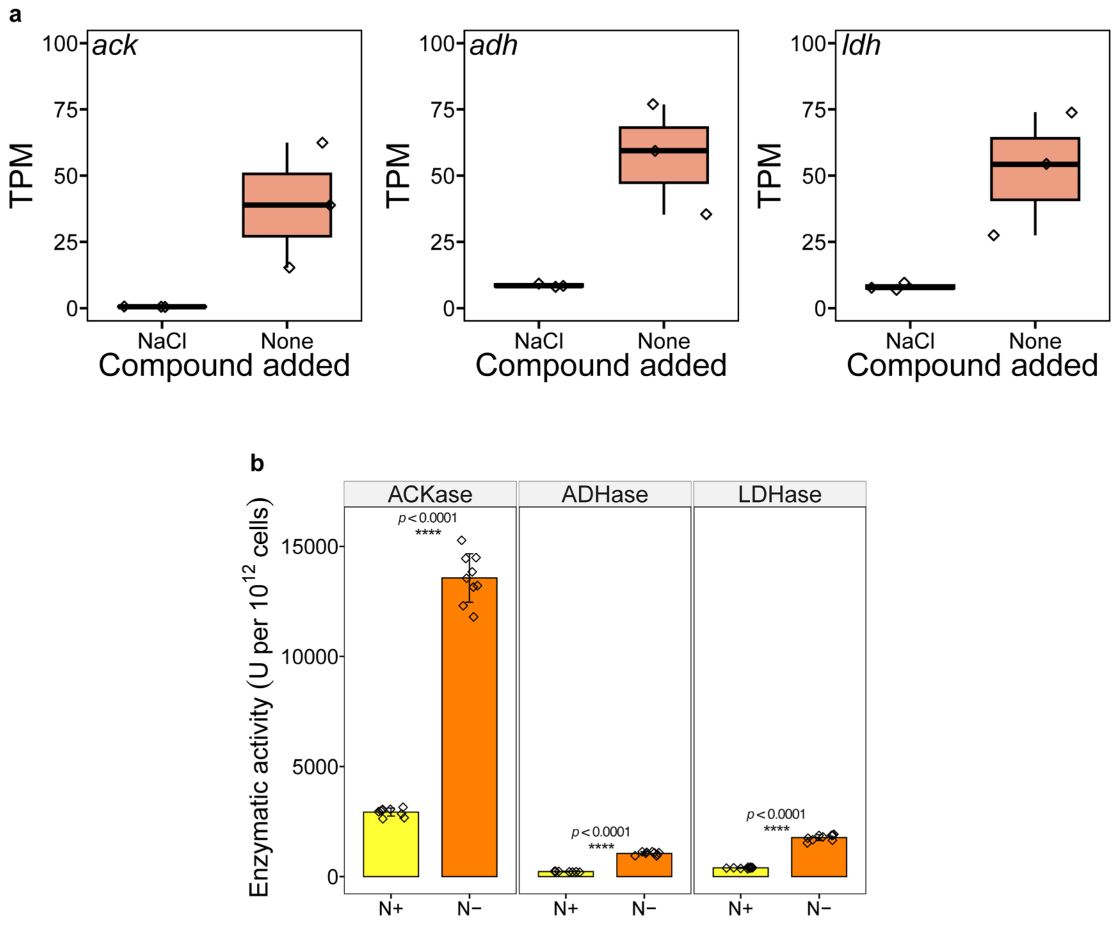


**Extended Data Fig. 6. Transcript analysis of fermentation involved genes and enzymatic activities of enzymes. a** *ack* encoding acetate kinase, *adh* encoding alcohol dehydrogenase, *ldh* encoding lactate dehydrogenase. All three genes were identified as significantly different expression genes between these two treatments ($p < 0.05$). Statistical significance of genes analysed was calculated from TPM (transcripts per million) pairwise comparisons of triplicate samples of NaCl deprivation treatment (None) compared to the NaCl addition treatment (NaCl). **b** Enzymatic activities of ACKase, ADHase and LDHase. “N+” indicates BG11 with NaCl (18 g/L) while “N-” denotes $BG11_0$ with NaCl (18 g/L). The graphs show the mean ± standard deviation (n = 9). Corresponding significance markers represent statistical significance calculated by Welch’s *t* test. Significance: ns (no significance), *($p < 0.05$), **($p < 0.01$), ***($p < 0.001$), ****($p < 0.0001$).

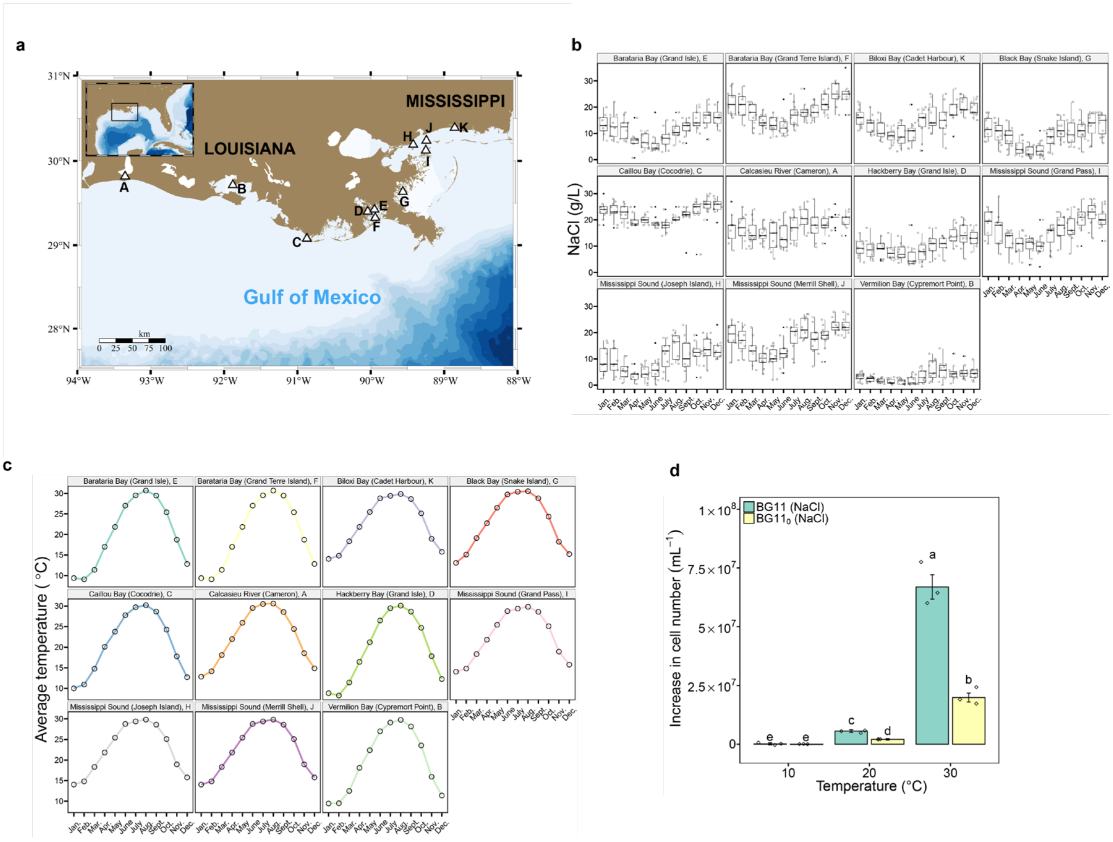


**Extended Data Fig. 7. Annual salinity and temperature dynamics of eleven coastal stations along the Texas coast of North America. a** Locations of stations. A, Calcasieu River (Cameron); B, Vermilion Bay (Cypremort Point); C, Caillou Bay (Cocodrie); D, Hackberry Bay (Grand Isle); E, Barataria Bay (Grand Isle); F, Barataria Bay (Grand Terre Island); G, Black Bay (Snake Island); H,

Mississippi Sound (Joseph Island); I, Mississippi Sound (Grand Pass); J, Mississippi Sound (Merrill Shell); K, Biloxi Bay (Cadet Harbour). **b** Annual salinity dynamics of eleven stations over the last decade. **c** Annual temperature dynamics of the surface water of the eleven stations. **d** Increase in cell number of populations grown in $BG11_0$ (NaCl) and BG11 (NaCl) at 10°C, 20°C and 30°C for seven days, respectively. The graph shows the mean ± standard deviation of the triplicate values. Different letters on each bar represent statistical significance ($p < 0.05$) calculated by ANOVA with Tukey's HSD post-hoc analysis across all populations tested.
